# Encephalitic Alphavirus Infection Induces PARP-1 Hyperactivation Mediated Energy Collapse in Motor Neurons

**DOI:** 10.64898/2026.01.23.701280

**Authors:** Rodney Eric Williams, Lisa Pieterse, Swara S. Patel, Mathew W. McLaren, Matthew J. Elrick, Diane E. Griffin

## Abstract

Motor neurons are highly vulnerable to metabolic stress, yet the mechanisms driving their degeneration during neurotropic alphavirus infections remain unclear. Venezuelan equine encephalitis virus (VEEV) causes motor neuron injury, but the intrinsic pathways underlying this susceptibility are not fully defined. Previous work suggests alphavirus-infected motor neurons may die through caspase-independent mechanisms. Here, we show that VEEV infection induces sustained activation of the DNA repair enzyme poly(ADP-ribose) polymerase-1 (PARP-1), leading to depletion of NAD^+^ and ATP in murine NSC34 motor neuron–like cells and human iPSC-derived motor neurons. These metabolic changes precede mitochondrial depolarization and cell death. Pharmacological inhibition or genetic reduction of PARP-1 partially restores NAD^+^ and ATP and improves cell survival, indicating that PARP-1 hyperactivation directly contributes to energetic collapse and intrinsic motor neuron death. These results identify PARP-1 as a key driver of energy failure during VEEV infection and a potential target to limit neuronal injury in neurotropic viral infections.

## Introduction

Alphaviruses are positive-sense, single-stranded, mosquito-borne RNA viruses of the *Togaviridae* family that can cause severe neurological disease in humans and animals (Lantz & Baxter, 2025; Woodson et al., 2025; Zacks & Paessler, 2010). They are broadly classified into Old World and New World lineages (Abdelnabi & Delang, 2020; Lantz & Baxter, 2025; VanderGiessen et al., 2025). New World alphaviruses, including Venezuelan (VEEV), Eastern, and Western equine encephalitis viruses, exhibit pronounced neurotropism and cause encephalitis and encephalomyelitis (Griffin, 2010; Lantz & Baxter, 2025; Weaver et al., 2004). VEEV is clinically significant due to its continued persistence in Central and South America and capacity to invade the central nervous system (CNS) and induce acute and chronic neurological sequelae or death (Guzman-Teran et al., 2020; Rivera et al., 2024; VanderGiessen et al., 2024; Woodson et al., 2025). Despite this clinical relevance and the considerable disease burden associated with alphavirus neuropathogenesis, there are currently no approved antiviral therapies targeting these viruses (Abdelnabi & Delang, 2020; Han et al., 2023; Ogorek & Golden, 2023; Stromberg et al., 2020).

Neurotropic alphavirus infections result in neuronal death in discrete regions of the CNS, including the cortex, hippocampus, brain stem, and spinal cord (Griffin, 2016; Potter et al., 2015). In severe infections, motor neuron injury can cause paralysis due to acute myelitis, often leading to long-term motor impairment (Kerr et al., 2002; Potter et al., 2015; Ronca et al., 2016). While apoptosis, necrosis, and inflammatory forms of cell death have been described in cortical and hippocampal neurons following alphavirus infection, the intrinsic cellular pathways that drive motor neuron death remain poorly defined (Darman et al., 2004; Havert et al., 2000; Nargi-Aizenman & Griffin, 2001; Nargi-Aizenman et al., 2004). Prior work from our laboratory and others has suggested that motor neuron degeneration may also involve extrinsic, non–cell autonomous mechanisms, including excitotoxic stress from surrounding cells (Darman et al., 2004; Nargi-Aizenman et al., 2004). However, it remains unclear how intrinsic pathways within motor neurons contribute directly to their vulnerability.

Motor neurons are among the most metabolically demanding cells (De Silva et al., 2022). Their long axonal projections impose substantial bioenergetic requirements to support axonal transport, synaptic activity, and structural maintenance (De Silva et al., 2022; Niven, 2016; Rajan & Fame, 2024; Vergara et al., 2019). Motor neurons rely on efficient energy metabolism to maintain ATP levels, calcium balance, and overall cellular homeostasis, but they have limited capacity to compensate for disruptions in energy supply or metabolic stress (De Silva et al., 2022). This energetic fragility is reflected in neurodegenerative disorders such as amyotrophic lateral sclerosis (ALS), where early deficits in metabolism and mitochondrial function drive disease progression (Tefera & Borges, 2017). The high metabolic demands of motor neurons suggest that disruptions in energy homeostasis may be a key contributor to their dysfunction and death during viral infection. Previous studies have reported mitochondrial dysfunction in neurons following VEEV infection (Keck et al., 2017), further supporting the role of metabolic stress in alphavirus neuropathogenesis.

Poly(ADP-ribose) polymerase-1 (PARP-1) is a nuclear enzyme that maintains DNA homeostasis by responding to cellular stress and catalyzing the synthesis of poly(ADP-ribose) (PAR) polymers using NAD^+^ as a substrate (Ray Chaudhuri & Nussenzweig, 2017). Under normal conditions, PARP-1 plays a critical role in DNA repair and genomic integrity. However, excessive PARP-1 activation can rapidly deplete NAD^+^ and reduce ATP availability, ultimately triggering mitochondrial depolarization and caspase-independent cell death, such as parthanatos (Fatokun et al., 2014). PARP-1 has also been implicated in antiviral responses, chromatin remodeling, and the regulation of inflammatory signaling during viral infection. Several viral infections, including herpesviruses, influenza virus, and HIV, induce robust PARP-1 activation and PAR accumulation, highlighting its dual roles as a DNA damage sensor and a potential modulator of viral replication (Bueno Murilo T. D. et al., 2013; Grady et al., 2012; Xia et al., 2020).

In alphavirus infection, prior studies in neuronal cells have documented PAR accumulation and suggested that PARP-1 hyperactivation may contribute to neuronal death through various pathways, including via depletion of NAD^+^ and ATP (Abraham et al., 2018; Nargi-Aizenman et al., 2002; Park & Griffin, 2009; Ubol et al., 1996). However, these studies have not been extended to motor neurons, which may exhibit distinct vulnerabilities due to their unique metabolic requirements and axonal architecture, and the role of PARP-1 mediated metabolic stress in neuronal cell vulnerability following infection remains incompletely defined.

Here, we hypothesize that VEEV infection triggers a PARP-1-mediated metabolic catastrophe in motor neurons. To test this, we utilized murine NSC34 motor-neuron-like cells as an established model for alphavirus-induced neuronal responses (Abraham et al., 2018; Burdeinick-Kerr & Griffin, 2005; Park & Griffin, 2009) and human iPSC-derived motor neurons (diMN) to capture human-specific physiological responses (NeuroLINCS Consortium et al., 2021). Specifically, we investigated whether viral-induced PARP-1 hyperactivation drives the depletion of NAD^+^ and ATP. Furthermore, we examined whether pharmacologic and genetic inhibition of PARP-1 or NAD^+^ supplementation could mitigate cellular metabolic collapse and improve neuronal survival, independent of viral replication.

## Results

### VEEV robustly infects motor neurons and induces cell death

To establish an *in vitro* model of VEEV neuropathogenesis in motor neurons, we first quantified viral replication in NSC34 cells and diMNs. Following infection with the attenuated VEEV-TC83 (VEEV) strain, plaque assays demonstrated efficient viral replication in both models, with titers increasing steadily between 6 and 36 hours post-infection (hpi) and reaching peak levels by 36 hpi before declining by 48 hpi (Figure 1A). Human diMN supported slightly higher peak titers than NSC34 cells. Cell death was evident after infection of both cell types as measured by Trypan blue assay, with the total number of surviving cells relative to baseline decreasing progressively over the course of infection (Figure 1B). By 6 hpi, the number of surviving cells had declined by approximately 25% and by 24 hpi survival was reduced by 45-55%. At 48 hpi, survival in diMN cultures was reduced by ∼75%, compared with ∼65% NSC34 cultures. Confocal imaging of VEEV-TC83-GFP infected cultures corroborated these findings, with widespread infection observable in infected cultures at 24 hpi (Figure 1C). Together, these data establish a robust *in vitro* motor neuron infection model, characterized by high viral burden and cell death.

**Figure 1.**
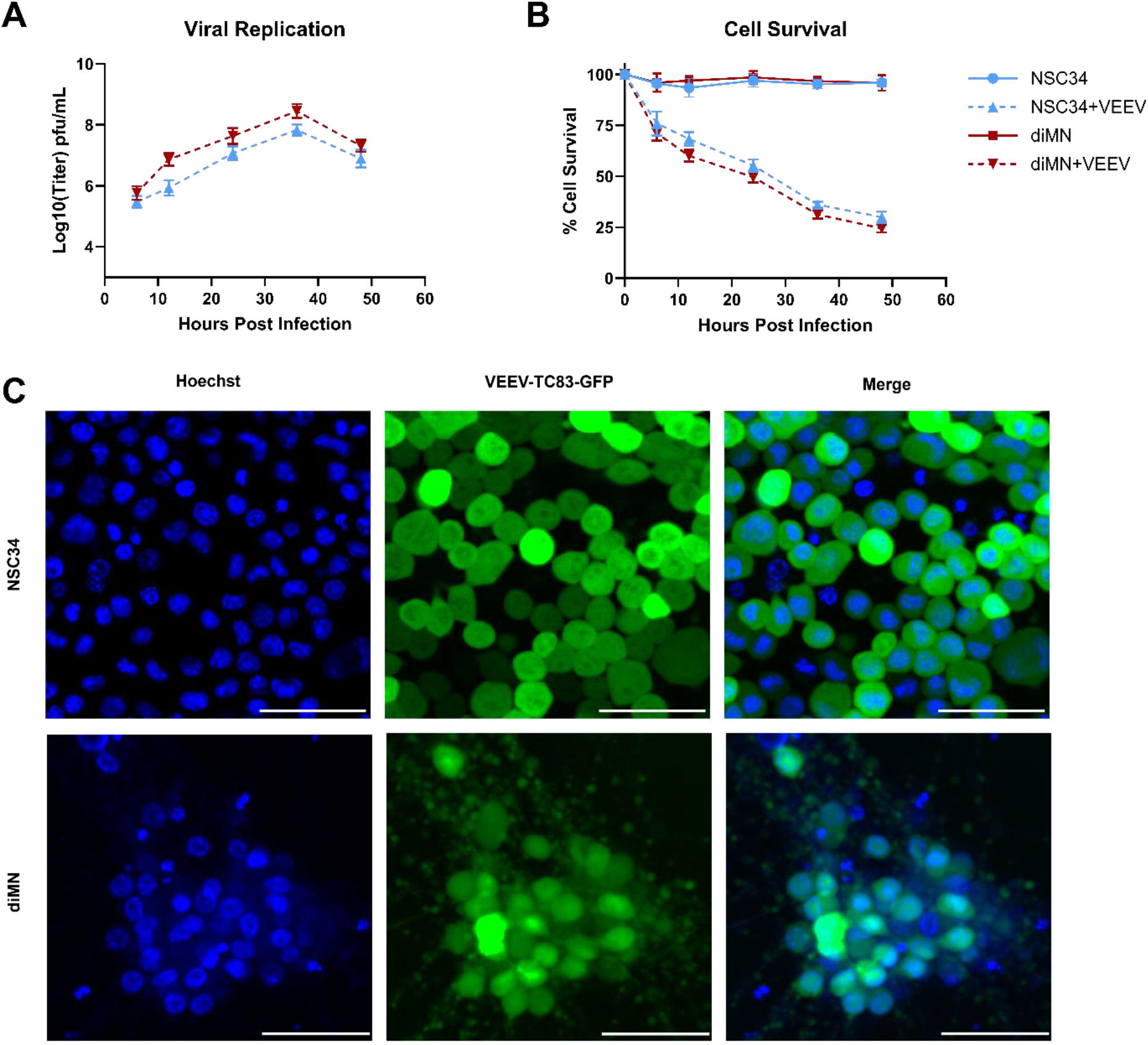
VEEV robustly infects motor neurons and induces progressive cell death. (**A**) Viral replication kinetics in NSC34 and diMN. Cells were infected with VEEV-TC83 (VEEV, MOI 1.0) and viral titers were quantified from supernatants at the indicated time points via plaque assay on Vero cells. Data represent mean ± SEM (PFU/mL) from three independent biological replicates. Both motor neuron models supported efficient viral replication, reaching peak titers by 36 hpi. (**B**) Cell survival in VEEV infected motor neurons. The number of surviving cells was quantified at 6, 12, 24, 36, and 48 hpi using the trypan blue exclusion assay. Data are presented as the percent of live cell relative to the total number of cells at 0 HPI (mean ± SEM). (**C**) Representative confocal microscopy images of NSC34 and diMN cultures at 24 hpi following infection with VEEV-TC83-GFP (green, MOI 1.0). Cellular nuclei were counterstained with Hoechst dye (blue). Widespread GFP expression demonstrates high infection efficiency across the motor neuron cultures. Scale bars = 50µm.

### VEEV induces sustained PARP-1 activation and depletion of NAD and ATP

To determine whether VEEV infection alters PARP-1 activity in motor neurons, we examined the accumulation of PAR following infection. Western blot analysis revealed PAR accumulation by 6 hpi in both infected NSC34 cells and diMN cultures, but not mock infected cultures, indicating early activation of PARP-1 (Figure 2A). PAR levels increased substantially by 12 hpi and remained elevated for the duration of the infection. Total PARP-1 protein levels remained relatively stable, suggesting that increased PAR accumulation reflects enzymatic hyperactivation rather than changes in PARP-1 expression.

**Figure 2.**
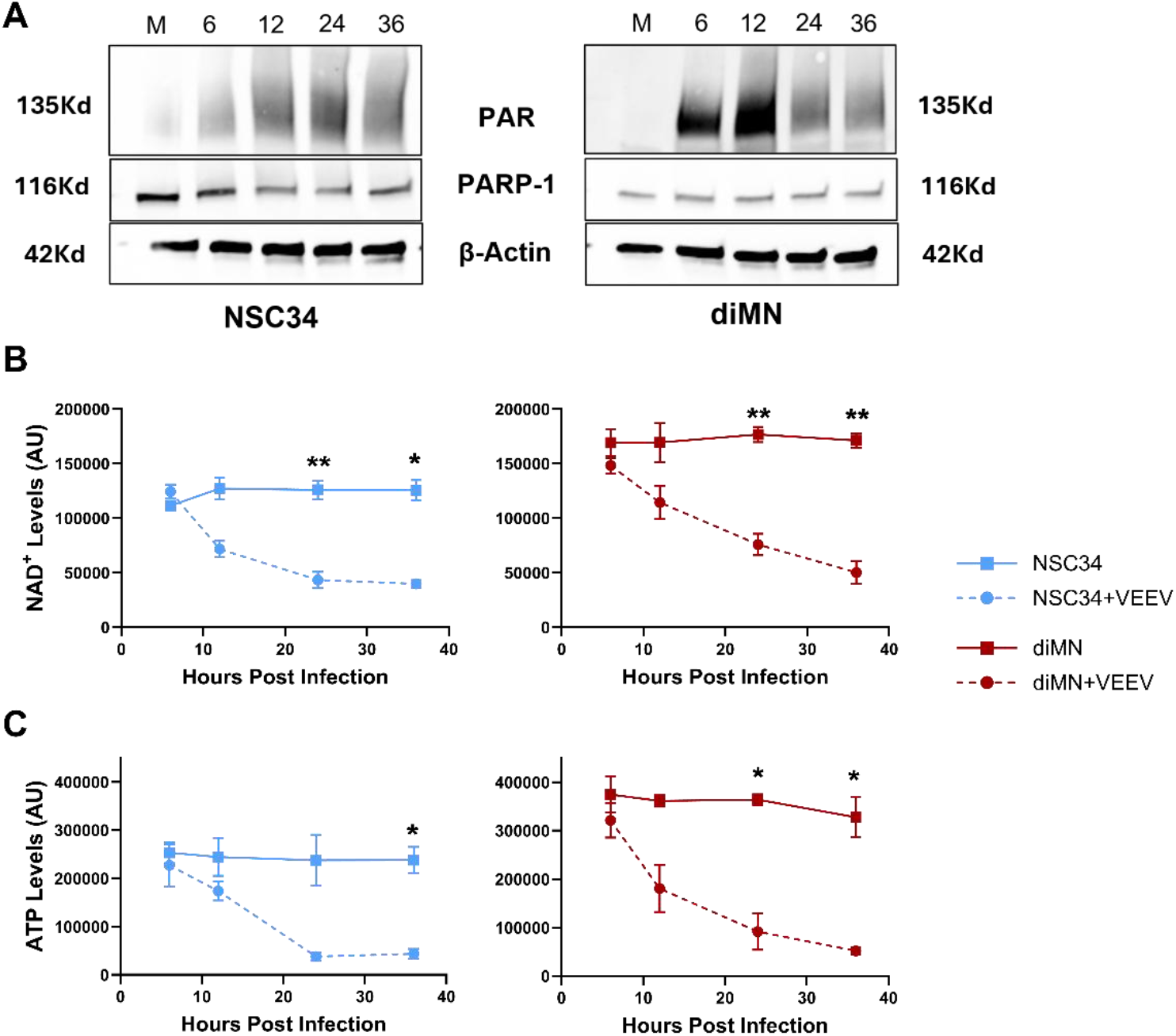
VEEV infection induces PARP-1 hyperactivation and depletion of NAD and ATP. (**A**) Representative Western blot analysis of PAR accumulation and PARP-1 expression in NSC34 and diMN cells at 6, 12, 24, and 36 hpi with VEEV (MOI 1.0). (**B**) NAD^+^ levels in diMN (red) and NSC34 (blue) cells infected with VEEV (MOI 1.0) across a 36-hour time course. Raw luminescent signals were normalized to live cell counts and are expressed as mean ± SEM luminescence per 10,000 cells from three independent experiments. Two-way ANOVA revealed a highly significant main effect of infection (p < 0.0001); asterisks indicate significance at specific time points determined by Sidak’s multiple comparisons post-hoc test. (**C**) ATP levels in diMN and NSC34 cells following VEEV infection. Data were normalized to live cell counts as described in (B). Main effect of infection was significant by Two-way ANOVA (p < 0.0001). Asterisks indicate results of Sidak’s multiple comparisons test at indicated hpi. * = P ≤ 0.05, ** = P ≤ 0.01.

Given that PARP-1 consumes NAD^+^ as a substrate when activated, we next assessed intracellular NAD^+^ levels across the same time course. NAD^+^ levels remained relatively stable at 6h hpi but declined sharply by 12 hpi, coinciding with the onset of robust PAR accumulation (Figure 2B). By 24 hpi, NAD^+^ levels were reduced nearly 3-fold when compared to mock-infected controls and remained markedly suppressed 36 hpi. These data indicate that sustained PARP-1 activation during VEEV infection is associated with rapid depletion of cellular NAD^+^ pools.

Because NAD^+^ availability is critical for cellular bioenergetics, we next assessed intracellular ATP levels following infection. Similar to NAD^+^ dynamics, ATP levels were largely preserved at 6 hpi but declined rapidly by 12 hpi in both motor neuron models (Figure 2C). By 24 hpi ATP levels had decreased 3-fold when compared to mock-infected controls, with levels in NSC34 cells reaching a nadir before rebounding slightly by 36 hpi. ATP levels in diMN cultures continued to progressively decline through 36 hpi. The temporal coupling of NAD^+^ depletion and ATP loss suggests that PARP-1-mediated NAD^+^ consumption contributes to downstream energetic collapse during VEEV infection.

Together these data demonstrate that VEEV infection induces early and sustained PARP-1 hyperactivation in motor neurons, followed by rapid depletion of NAD^+^ and ATP. Importantly PARP-1 hyperactivation precedes or coincides with metabolic failure, supporting a model in which dysregulated PARP-1 activity contributes to bioenergetic collapse during infection rather than arising solely as a secondary consequence of cell death.

### PARP-1 hyperactivation and NAD^+^/ATP depletion precede mitochondrial dysfunction and oxidative stress

To determine whether the metabolic collapse observed during VEEV infection was associated with downstream mitochondrial dysfunction, we next assessed changes in mitochondrial membrane potential (MMP) over the course of infection. MMP was measured using tetramethylrhodamine ethyl ester (TMRE) fluorescence in NSC34 cells and diMN at the same time points used for PARP-1 activation and metabolic analyses.

Despite early and robust PARP-1 activation and depletion of NAD^+^ and ATP, no significant loss of MMP was observed at 6 or 12 hpi in either motor neuron model (Figure 3A). A significant reduction in TMRE fluorescence first became evident at 24 hpi, indicating loss of mitochondrial membrane potential, and this decrease was sustained or further exacerbated by 36 hpi. These findings suggest that mitochondrial depolarization may occur downstream of early metabolic disruption rather than coinciding with the initial stages of PARP-1 activation.

**Figure 3.**
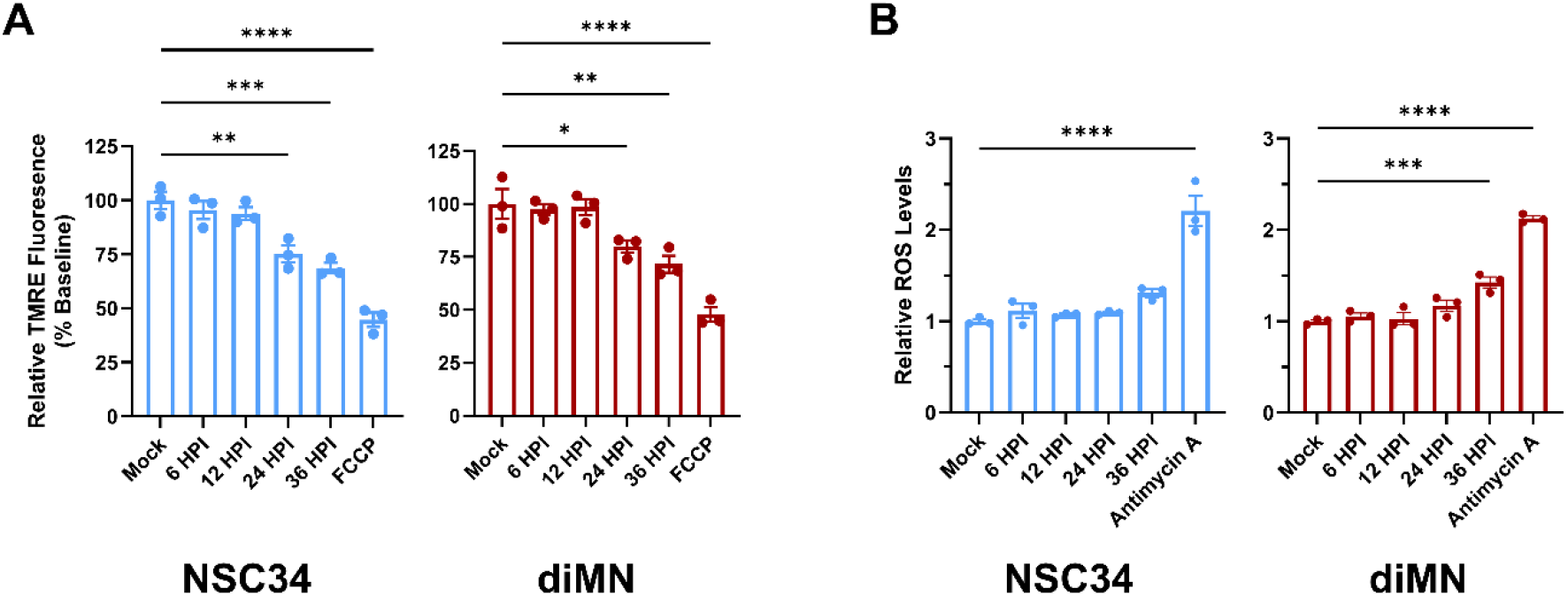
PARP-1 hyperactivation and NAD^+^/ATP depletion precede mitochondrial dysfunction and oxidative stress. (**A**) Mitochondrial membrane potential (MMP) was assessed in NSC34 and diMN cells from 6 to 36 hpi with VEEV (MOI 1.0) using TMRE (200nM) fluorescence. Data were normalized to total cell number and are expressed relative to uninfected Mock control at baseline. Carbonyl cyanide-p-(trifluoromethoxy)phenylhydrazone (FCCP, 20µM) was included as a positive control for mitochondrial depolarization, with addition occurring 10 minutes prior to fluorescent analysis. (**B**) Intracellular reactive oxygen species (ROS) levels were quantified over the same time course using DCFDA fluorescence. Signals were normalized to total live cell counts and are presented relative to the uninfected Mock control. Antimycin A (1µM) was used as a positive control for ROS induction, with addition occurring 15 minutes prior to fluorescent analysis. Statistical significance for (A) and (B) was determined by One-way ANOVA assuming a lognormal distribution with Dunnett’s multiple comparisons test, comparing each infected time point to the uninfected Mock control. Data represent mean ± SEM from three independent biological replicates.

Because mitochondrial dysfunction is often associated with increased oxidative stress, we next quantified intracellular reactive oxygen species (ROS) levels over the same time course. ROS levels remained comparable to mock-infected controls at early time points and showed only a slight increase at 24 hpi (Figure 3B). A noticeable elevation in ROS was not observed until 36 hpi with a significant increase in diMN and a similar trend in NSC34, temporally coinciding with advanced mitochondrial dysfunction and peak viral replication.

Together, these data demonstrate that PARP-1 hyperactivation and depletion of NAD^+^ and ATP occur prior to measurable mitochondrial depolarization and oxidative stress during VEEV infection. This temporal sequence supports a model in which early metabolic failure precedes and may contribute to subsequent mitochondrial dysfunction and secondary oxidative stress, rather than oxidative damage serving as the primary initiating event.

### PARP-1 inhibition and NAD^+^ supplementation partially restore cellular energy homeostasis and cell viability during VEEV infection

To determine whether PARP-1 hyperactivation directly contributes to metabolic dysfunction and reduced survival during VEEV infection, we next examined the effects of pharmacological and genetic inhibition of PARP-1 and NAD^+^ supplementation on PAR accumulation, viral replication, cellular energetics, and cell viability. NSC34 cells and human diMN were infected with VEEV and treated with the PARP inhibitor ABT-888 (PARPi), PARP-1–targeting siRNA (siPARP-1; NSC34 only), the NAD^+^ precursor nicotinamide riboside (NR), or combinations thereof, and outcomes were assessed primarily at 24 hpi.

Western blot analysis demonstrated robust PAR accumulation in infected cells at 24 hpi, which was markedly reduced by PARP inhibition or knockdown (Figure 4A). Treatment with ABT-888 substantially attenuated PAR accumulation in both NSC34 cells and diMN, while siRNA-mediated knockdown of PARP-1 in NSC34 cells similarly reduced PAR signal relative to infected controls.

**Figure 4.**
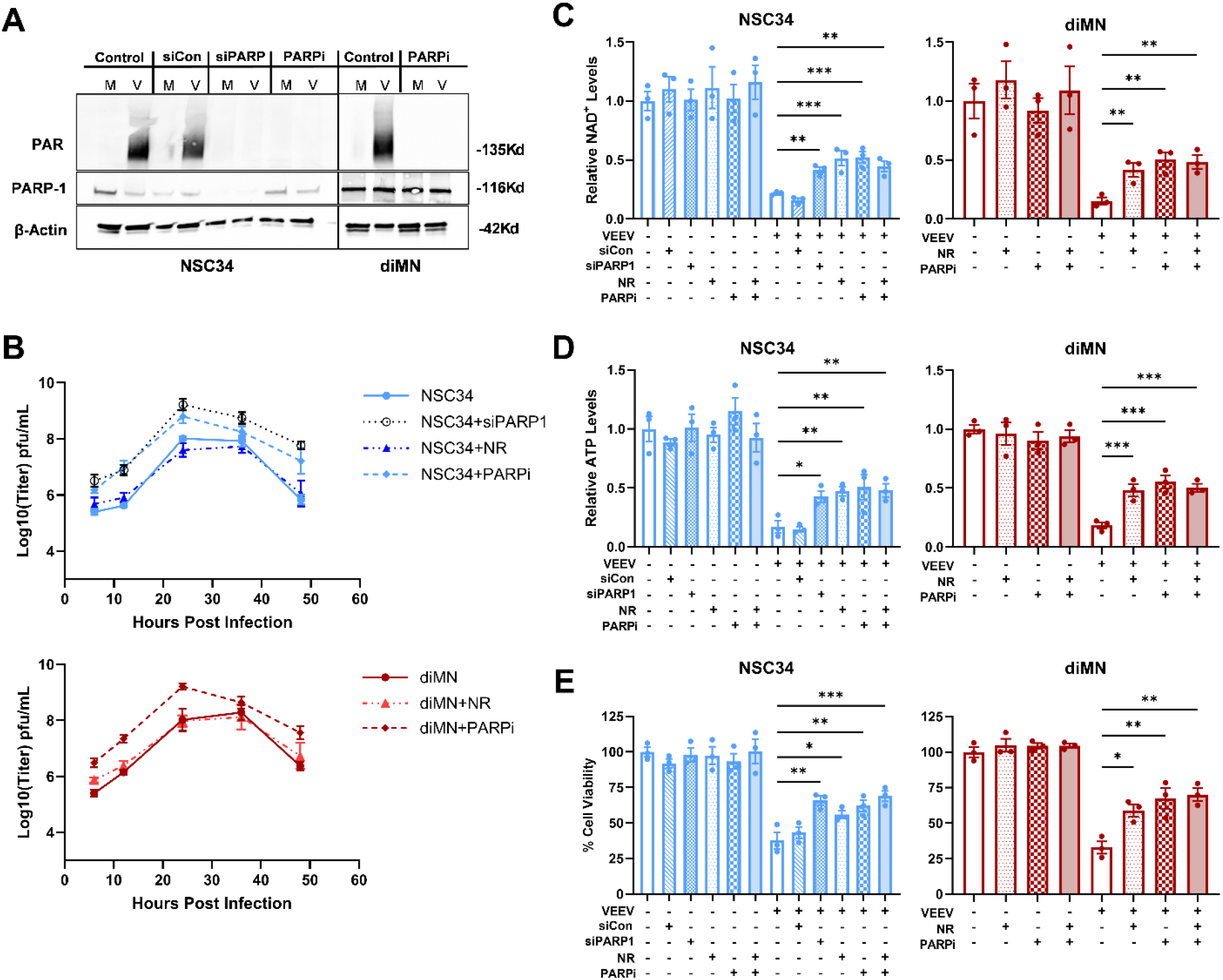
PARP-1 inhibition and NAD^+^ supplementation partially restore cellular energy homeostasis and viability during VEEV infection. (**A**) Representative western blot analysis of PAR accumulation and PARP-1 expression at 24 hpi with VEEV (MOI 1.0). NSC34 cells were transfected with either PARP1 siRNA (siPARP1; 20 pmol) or scrambled control siRNA (siCon; 20 pmol) 48 hours prior to infection. Pharmacological inhibition was achieved using ABT-888 (PARPi; 20µM) initiated 1 hour prior to infection and maintained throughout the experiment. (**B**) VEEV replication kinetics in NSC34 and diMN cells (MOI 1.0) following PARP-1 manipulation or nicotinamide riboside (NR; 500 µM) supplementation. To maintain effective concentrations, a 250µM NR spike was added to the infectious media at 12 hpi. Viral titers were quantified by plaque assay on Vero cells from supernatants collected between 6 and 48 hpi. Data represent mean ± SEM from at least three independent biological replicates. PARP-1 inhibition (siPARP1 and PARPi) resulted in an approximate one-log increase in viral titers between 12 and 24 hpi. **(C–E)** Metabolic and viability rescue at 24 hpi. (C) Intracellular NAD^+^ (NAD/NADH-Glo) and (D)ATP levels (Luminescent ATP Detection) were quantified following treatment with siCon, siPARP1, PARPi, or NR as described above. All metabolic values were normalized to the number of viable cells in matched wells on the same plate to account for infection-induced cell loss. (E) Cell viability was assessed via Alamar Blue fluorescence. All values are expressed relative to mean of uninfected Mock controls. Statistical significance was determined by ordinary one-way ANOVA assuming a lognormal distribution; infected-treated groups were compared to the infected-untreated (or siCon) control via Dunnett’s multiple comparisons test. Mock and treatment-only controls are shown for reference. Data represent mean ± SEM., * = P ≤ 0.05, ** = P ≤ 0.01, *** = P ≤ 0.001

We next assessed the impact of PARP-1 modulation and NAD^+^ supplementation on viral replication kinetics. Notably, pharmacological or genetic inhibition of PARP-1 resulted in increased viral titers in both NSC34 and diMN (Figure 4B). In contrast, NR supplementation did not alter viral replication in either cell type. These results indicate that the metabolic rescue provided by PARP-1 inhibition occurs independently of viral burden, as these interventions preserve cellular viability despite high levels of ongoing viral replication. Furthermore, these data demonstrate that PARP-1 inhibition does not exert a generalized antiviral effect but rather acts specifically to prevent the metabolic collapse downstream of infection.

We then examined whether inhibition of PARP-1 activity or NAD^+^ supplementation could rescue infection-associated depletion of NAD^+^ and ATP. In both NSC34 cells and diMN, VEEV infection resulted in substantial reductions in intracellular NAD^+^ levels at 24 hpi relative to mock-infected controls (Figure 4C). Treatment with PARPi, siPARP-1 (NSC34), NR, or combined PARPi and NR significantly restored NAD^+^ levels. Similar trends were observed for ATP levels (Figure 4D), with all treatments coinciding with a significant partial restoration of ATP concentrations. Together, these findings indicate that both suppression of PARP-1 activity and replenishment of NAD^+^ pools can mitigate infection-induced energy depletion.

Finally, we assessed whether modulation of PARPi activity or NAD^+^ supplementation translated into improved cellular viability. At 24 hpi, VEEV infection caused a pronounced reduction in cell viability as measured by Alamar blue reduction in both NSC34 cells and diMN (Figure 4E). Treatment with PARPi, siPARP-1 (NSC34), NR, or combined PARPi and NR significantly increased viability relative to infected untreated controls, with approximately 2–2.5-fold improvements observed across treatment groups. PARP-1 inhibition and combined PARPi and NR treatment produced the most robust increases in viability in both cell types, while NR alone also significantly improved viability without enhancing viral replication. These results indicate that modulation of PARP-1 activity and NAD^+^ availability partially restores metabolic function and improves neuronal survival during VEEV infection.

Collectively, these data demonstrate that excessive PARP-1 activity contributes to metabolic collapse and cell death in infected motor neurons. While inhibition of PARP-1 enhances viral replication, restoration of NAD^+^ and ATP levels through either PARP-1 suppression or NAD^+^ supplementation improves metabolic function and cell survival, supporting a model in which PARP-1–driven energy depletion is a key determinant of motor neuron vulnerability during alphavirus infection.

These results demonstrate that VEEV infection of motor neurons is associated with early and sustained PARP-1 activation, followed by depletion of intracellular NAD^+^ and ATP, mitochondrial dysfunction, and reduced cellular viability. Modulation of PARP-1 activity or NAD^+^ availability partially preserved metabolic parameters and improved survival despite continued viral replication. Together, these findings support a model in which PARP-1– associated metabolic stress contributes to motor neuron vulnerability during alphavirus infection.

## Discussion

In this study, we show that PARP-1 hyperactivation by VEEV infection leads to NAD and ATP depletion and cell death in motor neurons. Inhibition of PARP-1 or supplementation of NAD preserves metabolic function and improves survival of motor neurons during VEEV infection, despite permitting continued viral replication. These findings support a model in which PARP-1 activation contributes directly to host cell metabolic failure and neuronal injury, rather than serving solely as a downstream marker of cellular stress. Importantly, this effect was observed in both murine motor neuron–like cells and human iPSC-derived motor neurons, indicating that PARP-1–driven metabolic vulnerability is conserved across systems.

Our data indicate that PARP-1 activation precedes measurable declines in intracellular NAD^+^ and ATP levels, followed by loss of mitochondrial membrane potential and reduced cell viability. While mitochondrial dysfunction has been reported previously during alphavirus infection (Keck et al., 2017), our results suggest that disruption of cellular energy balance doesn’t exclusively follow from mitochondrial failure. Excessive PARP-1 activity consumes NAD^+^ and limits ATP availability at earlier timepoints than mitochondrial membrane depolarization or ROS accumulation. Motor neurons appear poorly equipped to buffer these metabolic disruptions. This is consistent with the high energetic demands of motor neurons and their limited capacity to compensate for sustained energy perturbations (De Silva et al., 2022; Keck et al., 2017; Niven, 2016; Rajan & Fame, 2024; Rumpf et al., 2023). Together, these findings place PARP-1–mediated NAD^+^ depletion as a proximal trigger for metabolic collapse. While multiple independent pathways of metabolic compromise likely converge during infection, our data suggest that PARP-1 activity represents an early event in the cascade.

While previous work noted PARP-1 activation in various neuronal lines (Abraham et al., 2018; Nargi-Aizenman et. al, 2002; Park & Griffin, 2009) and speculated on the role of energy loss (Ubol et al., 1996), our study establishes a direct causal relationship. By integrating pharmacologic inhibition, genetic knockdown, and metabolic supplementation, we demonstrate that PARP-1 activity is a primary driver of, rather than a secondary marker for, energetic failure and death in motor neurons.

Motor neuron death during neurotropic viral infection has also been partially attributed to extrinsic, non–cell autonomous mechanisms, including excitotoxic stress mediated by surrounding cells (Darman et al., 2004). Prior studies from our laboratory and others support an important role for these pathways, particularly in vivo (Burdeinick-Kerr & Griffin, 2005; Darman et al., 2004; Fannjiang et al., 2003; Greene et al., 2008; Nargi-Aizenman et al., 2004). The present findings do not contradict this model but instead indicate that intrinsic metabolic stress within motor neurons can independently drive cell death. PARP-1–dependent energy depletion may therefore act as a permissive or amplifying factor that lowers the threshold for motor neuron death in the context of additional extrinsic insults.

An important observation from this study is the dissociation between neuronal survival and viral replication. Both pharmacologic inhibition and siRNA-mediated reduction of PARP-1 activity moderately increased viral replication, consistent with prior evidence that PARP family members contribute to antiviral restriction (Malgras et al., 2021). This highlights a functional tradeoff in which PARP-1 activity supports antiviral defense but compromises host cell metabolic integrity. In contrast, supplementation with nicotinamide riboside restored NAD^+^ and ATP levels and improved cell viability without substantially enhancing viral replication, suggesting that metabolic support may be able to mitigate neuronal injury without disrupting antiviral control.

The use of human iPSC-derived motor neurons provides additional insight into the relevance of these mechanisms in a human cellular context. Human motor neurons recapitulated key features observed in murine cells, including PAR accumulation, metabolic depletion, and rescue with PARP inhibition or NAD^+^ supplementation. Given species-specific differences in innate immune signaling and neuronal metabolism (Hader et al., 2023; Iwata & Vanderhaeghen, 2024), these results strengthen the translational significance of PARP-1–mediated metabolic vulnerability in motor neurons.

Several limitations should be noted. These experiments were conducted in simplified *in vitro* systems and do not capture the contributions of glial cells, inflammatory mediators, or network-level activity that influence motor neuron survival *in vivo*. In addition, the upstream signals responsible for triggering PARP-1 hyperactivation during VEEV infection are still not completely understood. Viral replication–associated stress or innate immune signaling may each contribute and warrant further investigation (Sharma & Knollmann-Ritschel, 2019) . Finally, while NAD^+^ depletion was reproducibly observed, the extent to which ATP loss is directly driven by NAD^+^ consumption versus secondary effects of cellular stress remains to be fully defined.

In conclusion, our findings identify PARP-1 hyperactivation as a key mediator of metabolic failure and intrinsic motor neuron vulnerability during alphavirus infection. By directly linking PARP-1 activity to depletion of NAD^+^ and ATP and to reduced neuronal survival, this work clarifies a mechanism that complements existing models of extrinsic motor neuron injury. More broadly, these results emphasize the importance of metabolic homeostasis in determining neuronal outcome during viral infection and suggest that strategies aimed at preserving cellular energy balance may provide neuroprotection without fully compromising antiviral defense.

## Materials and Methods

### Viruses and Cell Lines

Venezuelan equine encephalitis virus (VEEV) vaccine strain TC-83 and the reporter virus TC-83–GFP were used in this study. VEEV TC-83 was originally provided by Dr. Ilya Frolov (University of Alabama at Birmingham), and TC-83–GFP was provided by Dr. Michael Diamond (Washington University in St. Louis) (Cain et al., 2017). Viral stocks were propagated in BHK-21 cells as previously described (Abraham et al., 2018). Briefly, cells were infected at a multiplicity of infection (MOI) of 0.1 and virus-containing supernatants were harvested 48 h post-infection, clarified by centrifugation, aliquoted, and stored at −80°C. Viral titers were determined by standard plaque assay on Vero cells (Abraham et al., 2018).

NSC34 motor neuron–like cells (a hybrid of mouse N18TG2 neuroblastoma cells and mouse spinal motor neurons) were originally a gift from Dr. Neil Cashman (University of Toronto). Cells were maintained at 37°C in DMEM supplemented with 10% fetal bovine serum, 1× penicillin–streptomycin, and 2 mM L-glutamine as previously described.

Direct-induced motor neurons (diMN) were generated from human iPSCs following an established differentiation protocol (NeuroLINCS Consortium et al., 2021). iPSC lines CS88, CS0YX7, CS8VTR were obtained from the Cedars-Sinai Biomanufacturing Center repository. Cells were maintained on UltiMatrix-coated 6-well plates in Essential 8 Flex media (Thermo Fisher), with differentiated cells manually removed, and passaged as colonies using ReLeSR (STEMCELL Technologies). Briefly, iPSCs were grown to ∼35% confluency and cultured in stage 1 media for 6 days to generate neuroepithelial cells. On day 6, cells were dissociated with Accutase, collected, and either cryopreserved or replated for further differentiation.

Thawed day-6 neuroepithelial cells were plated on UltiMatrix-coated plates in stage 2 media with 20 µM ROCK inhibitor Y-27632. Cells were fed daily with stage 2 media without ROCK inhibitor from days 7–12 to produce motor neuron precursors. On day 12, precursors were dissociated with 0.05% trypsin, replated into the experimental format. From day 12 onward, cells were cultured in stage 3 media to support terminal motor neuron maturation. Cultures contained ∼80% Tuj1+ neurons, of which roughly half were Islet-1+ mature motor neurons and half Nkx6.1+ immature motor neurons. diMN were used between Day 28–35 post-induction. For imaging experiments, cells were plated onto Ultimatrix-coated 24- or 96-well glass-bottom plates (Cellvis) at 2.5×105 cells/well and 5×104 cells/well, respectively. For biochemical assays, cells were maintained in 6-well plates throughout Stage 2–3. The three diMN cell lines served as independent biological replicates in all experiments, unless otherwise noted.

### Viral Infection of Motor Neurons

NSC34 and diMN cultures were infected with VEEV TC83 or TC83–GFP at an MOI of 1 unless otherwise indicated. Virus was diluted in serum-reduced medium (DMEM with 2% FBS for NSC34 or S3 medium for diMN). Cells were inoculated for 1 h at 37°C with gentle rocking every 15 min. After inoculation, viral inoculum was removed and cells were washed once with phosphate-buffered saline (PBS; pH 7.4) and treated with fresh culture media. Mock-infected controls received equivalent medium without virus. For time-course experiments, samples were collected at 6, 12, 24, 36, or 48 h post-infection (hpi), as indicated.

### Protein Extraction and Quantification

At designated timepoints post-infection, cells were washed with cold PBS and lysed in RIPA buffer supplemented with protease and phosphatase inhibitors. Lysates were clarified by centrifugation (10,000× g, 5 min, 4°C). Protein concentration was quantified by DC protein assay (Bio-Rad #5000111) according to the manufacturer’s instructions.

For Western blotting, 10µg of total protein was resolved per lane by SDS–PAGE gel electrophoresis and transferred to nitrocellulose membranes. Membranes were blocked in 5% BSA in TBST and incubated with primary antibodies against PAR (1:3000) (Kang et al., 2025)(gift from the lab of Ted and Valina Dawson at Johns Hopkins University), PARP-1 (1:1500)(Cell Signaling Technologies), or β-actin(1:5000) (Cell Signaling Technologies) at 4°C overnight. HRP-conjugated secondary antibodies (1:2000) were applied for 1 h at room temperature, and signal was developed using ECL prime chemiluminescent substrate (Cytiva #RPN2232) and imaged using the Bio-Rad Quantity-One imager and software.

### Quantification of Energy Substrates

Intracellular NAD^+^ and ATP levels were quantified using commercial luminescence-based assays. NAD^+^ content was measured using the NAD/NADH-Glo assay (Promega) following the manufacturer’s protocol. ATP levels were determined using the Luminescent ATP detection Kit (Abcam). Briefly, cells were lysed directly in assay buffer, incubated for 10 min, and luminescence was measured using a plate reader (Tecan Spark). All values were normalized to number of viable cells in matched wells on the same plate at the same time points.

### TMRE Mitochondrial Membrane Potential Assay

Mitochondrial membrane potential was assessed using a tetramethylrhodamine ethyl ester (TMRE) based assay (Abcam #ab113852). Cells were incubated with 200nM TMRE in culture medium for 30 min at 37°C. After incubation, cells were washed gently with warm PBS and quantified using a Tecan Spark high-throughput plate reader. TMRE signal intensity was normalized to total cell number and presented relative to mean of mock-infected controls at baseline. Carbonyl cyanide-p-(trifluoromethoxy)phenylhydrazone (FCCP) (20µM) was used as a depolarizing control.

### Reactive Active Oxygen Species Assay

Reactive oxygen species levels were measured using the Abcam DCFDA / H2DCFDA - Cellular ROS Assay Kit (#ab113851) according to the manufactures instructions and measured using a Tecan Spark microplate reader. Antimycin A was used as a positive control at 1µM for 15 minutes. Levels were normalized to the total number of live cells and presented relative to the mock untreated control at baseline.

### PARP-1 Inhibition and NAD^+^ Precursor Supplementation

#### ABT-888 (Veliparib)

PARP-1 activity was inhibited using ABT-888 (Veliparib) (Stemcell Technologies #100-1170). A 10mM stock was prepared in DMSO and stored at −20 °C protected from light. Working concentrations were prepared fresh by dilution into culture medium. Cells received 20µM ABT-888, added 1 h prior to infection, and maintained for the duration of the experiment unless otherwise indicated. Vehicle controls were treated with equivalent volumes of DMSO (<0.2% final).

### Nicotinamide Riboside (NR) Supplementation

To support NAD^+^ pools, cultures were treated with nicotinamide riboside (NR; Cayman chemicals #23132). NR free-base powder was dissolved fresh in sterile water to generate a 500mM stock, aliquoted, and stored at −80 °C. A 500µM working concentration was added directly to cultures 1 h pre-infection, during infection, and immediately post-infection. A 250µM concentration working stock was added as a 10x spike to infectious media at 12 hpi. Vehicle controls received medium alone.

### PARP-1 siRNA Knockdown

PARP-1 expression was reduced using siRNA targeting mouse PARP1 (Santa Cruz Biotechnology #sc-29438). A non-targeting scrambled siRNA (Santa Cruz, #sc-37007) served as a control. NSC34 cells were seeded at ∼60% confluence and transfected with 20pmol siRNA using Lipofectamine RNAiMAX (Invitrogen #13778) in Opti-MEM. After 24 h, medium was replaced with complete growth medium. Knockdown efficiency was assessed by Western blot 48 h post-transfection, and experiments were initiated at this time. ABT-888 was not used in siRNA experiments unless stated.

### Cell Survival and Viability Assays

Cell survival was quantified by trypan blue exclusion. Cells were harvested at the indicated timepoints with trypsin, mixed 1:1 with 0.4% trypan blue (Gibco #15250061), and counted using an Invitrogen Countess 3 automated cell counter. Percent survival was calculated as the ratio of live cells at each time point to the baseline number of cells at 0HPI.

Cell viability was measured using the resazurin-based Alamar Blue assay (Invitrogen #A50100). Cells were incubated with Alamar Blue reagent (10% v/v in media) for 4 h at 37°C, and fluorescence was measured at excitation/emission 560/590 nm using a Tecan Spark multiwell plate reader. Results were normalized to mock-infected controls.

### High-Content and Fluorescence Imaging

Cells were imaged using a Molecular Devices ImageXpress Micro Confocal platform equipped with 4×, 10×, 20×, and 40× objectives. For live imaging, cells were maintained at 37°C and 5% CO_2_ using an environmental chamber. GFP viral reporter signal and Hoechst, were captured with appropriate excitation/emission filter sets. Image analysis was performed using FIJI/ImageJ.

## Statistical Analysis

All experiments were performed in biological triplicate. Data are presented as mean ± SEM. Statistical analyses were performed using GraphPad Prism version 10.6. Comparisons between two groups were made using unpaired two-tailed Student’s t-tests. Multiple comparisons were performed using one-way or two-way ANOVA with appropriate post-hoc test unless otherwise stated. A p value < 0.05 was considered statistically significant.

## Author Contributions

Conceptualization: R.E.W. and D.E.G.; Investigation: R.E.W., L.P., and S.S.P.; Resources: M.W.M., M.J.E.; Formal Analysis: R.E.W.; Visualization: R.E.W.; Writing-original draft: R.E.W.; Writing-review & editing: R.E.W. and M.J.E.; Supervision: R.E.W., M.J.E. and D.E.G.; Funding Acquisition: D.E.G.

## Acknowledgements

This work is dedicated to the late Dr. Diane E. Griffin, who passed away on October 28, 2024. This work was supported by NIH Grants R01AI182066 and R56AI137264 (D.E.G.) and 1K08NS124989 and R01NS143998 (M.J.E.). S.S.P. was a participant in the Johns Hopkins Careers in Science and Medicine Initiative. We thank Aanishaa Jhaldiyal, Bong Gu Kang, and Rachy Abraham for helpful expertise on PARP-1 and PAR experiments and supply of related reagents. Figure 5 was created using BioRender.com.

**Figure 5.**
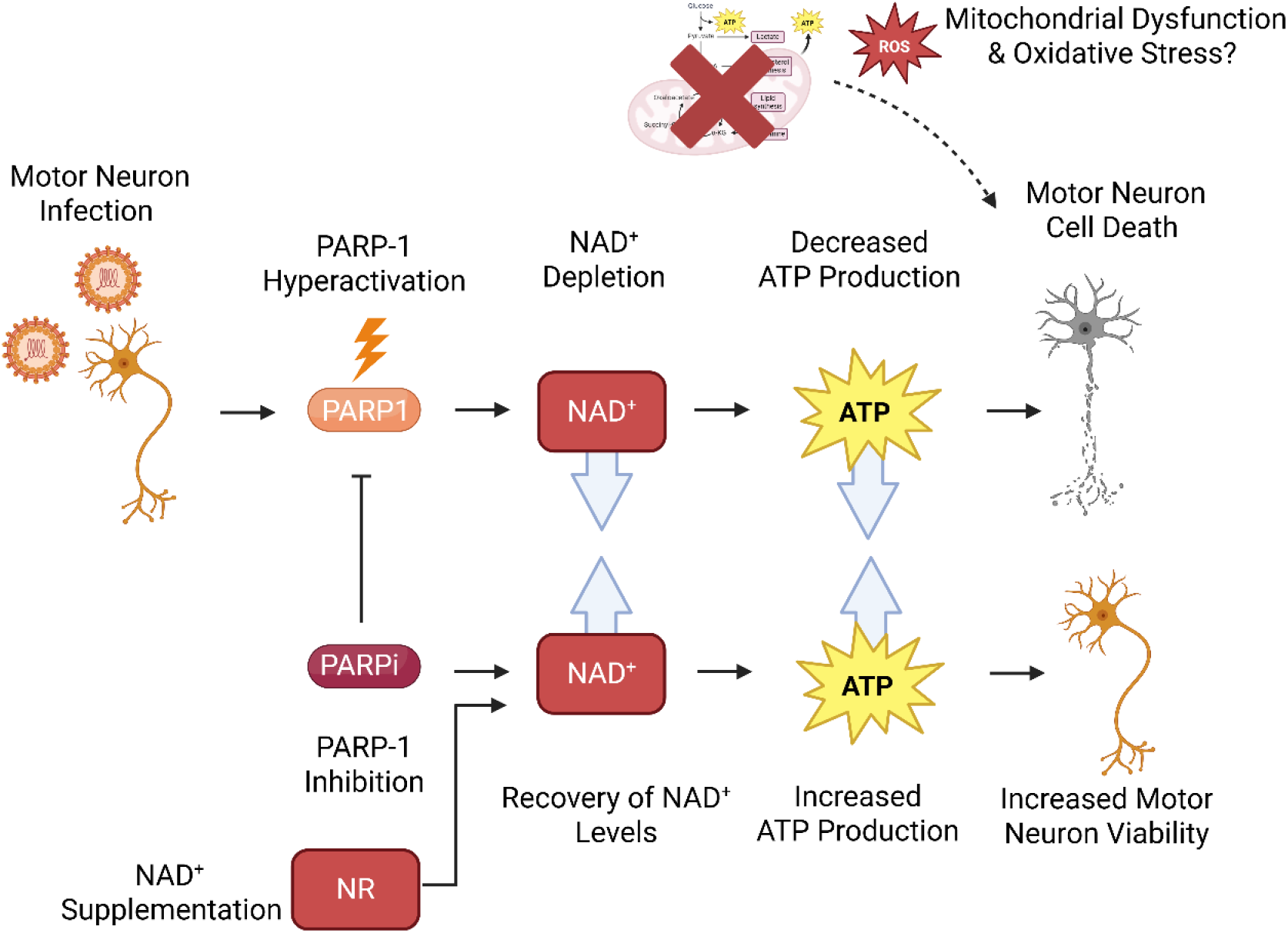
Illustrative model of VEEV-induced PARP-1 hyperactivation leading to metabolic collapse and neuronal death.

## Data Availability

All key data is directly shared within this manuscript. Any other data not shared in this manuscript will be made available upon request to the authors.

